# Balance between promiscuity and specificity in phage λ host range

**DOI:** 10.1101/2020.06.25.171868

**Authors:** Bryan Andrews, Stanley Fields

## Abstract

As hosts acquire resistance to viruses, viruses must overcome that resistance to re-establish infectivity, or go extinct. Despite the significant hurdles associated with adapting to a resistant host, viruses are evolutionarily successful and maintain stable coevolutionary relationships with their hosts. To investigate the factors underlying how pathogens adapt to their hosts, we performed a deep mutational scan of the region of the λ tail fiber tip protein that mediates contact with the λ host, *E. coli*. Phages harboring amino acid substitutions were subjected to selection for infectivity on wild type *E. coli*, revealing a highly restrictive fitness landscape, in which most substitutions completely abrogate function. By comparing this lack of mutational tolerance to evolutionary diversity, we highlight a set of mutationally intolerant and diverse positions associated with host range expansion. Imposing selection for infectivity on three λ-resistant hosts, each harboring a different missense mutation in the λ receptor, reveals hundreds of adaptive variants in λ. We distinguish λ variants that confer promiscuity, a general ability to overcome host resistance, from those that drive host-specific infectivity. Both processes may be important in driving adaptation to a novel host.

## Introduction

Viruses and their hosts engage in an evolutionary battle: mutations in the virus that increase infectivity come at a cost to bacterial growth, and mutations in the host that confer resistance to viruses come at a cost to the virus. In phages, as in other viruses, much of this battle is centered on the binding relationship between the host receptor and the viral protein that contacts it. Despite the adaptability of viruses to resistant hosts, viruses face significant evolutionary hurdles that their hosts do not (Fig. 1). First, random mutations are much more likely to disrupt an existing binding relationship than to generate a new one. Thus, viruses must survey a much broader sequence space than their hosts do to compete in an evolutionary arms race. Second, many receptors are readily lost in the host, but are essential to the virus. Some viruses, including λ, switch receptors following loss of their canonical receptor, although it is unclear if this is a common response [1, 2, 3]. Third, host populations are polymorphic, such that selection for viral resistance may give rise to multiple sub-clades with distinct resistance mechanisms. Overcoming any single resistance mechanism may not be sufficient for a virus to reestablish infectivity at the population level. Given these evolutionary hurdles, there remains the need to explain why viruses are so successful over evolutionary time scales [4, 5, 6]. In humans and other long-lived organisms, differences in relative generation times and mutation rates likely play a role [7]. However, phages are also extremely successful predators despite the short generation times and rapid adaptation of their bacterial hosts, which frequently outcompete phages and drive them to extinction in co-culture [6, 8].

**Figure 1:**
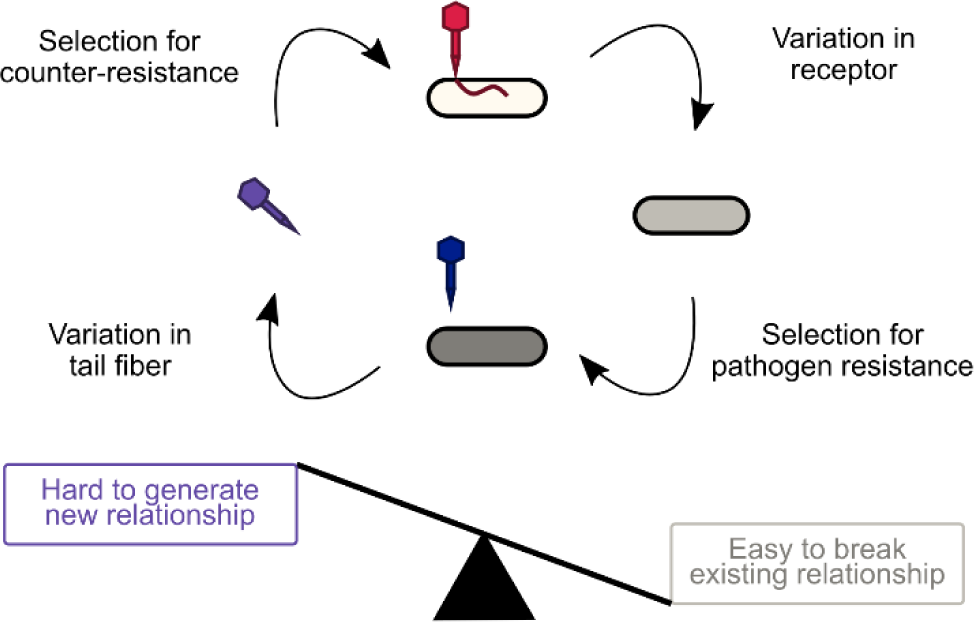
Phage–bacteria coevolution is fundamentally weighted in favor of bacteria. Phages and bacteria are presumed to coevolve stably and indefinitely in the wild, but theoretical considerations and experimental co-culture systems mostly predict long-term dominance of bacteria. Explaining the success of phages over evolutionary timescales is further hampered the limited mechanistic understanding of how phages adapt to a new and/or resistant host.

Phages such as λ serve as a useful model for analyzing host-pathogen coevolution [6, 8, 9]. Co-culture of λ with its *E. coli* host has resulted in the isolation of λ-resistant *E. coli* strains, which typically harbor null mutations of a maltose porin, LamB, that serves as the λ receptor [10, 11, 12]. Selection for both λ-resistance and maltose uptake has revealed LamB missense variants that disrupt λ binding [13]. λ variants can then arise that can re-establish infection on these previously resistant strains [13, 14]. When co-cultured with *E. coli* not expressing LamB, λ strains have been isolated that switch to use the non-canonical receptor OmpF, a LamB paralog [3].

Phage tail fibers, which bind to the host, evolve extremely quickly due to strong selective pressures and dedicated diversification mechanisms [2, 15, 16]. The λ tail fiber J protein consists of 1,132 amino acids, with high conservation across its N-terminal ∼980 amino acids and extreme diversification across its ∼150 amino acid C-terminal domain, which contacts the receptor [12, 14]. As examples, the J proteins from two recently isolated phages, lambda_2H10 and lambda_2G7b, are each >97% identical to λ across the N-terminal 982 amino acids, but only 40-60% identical across the C-terminal ∼150 residues [18]. The isolates are similarly diverged from each other and have unique host ranges. Although the structure of J or homologous tail fiber proteins has not been solved, tail fibers such as that of T4 form highly intertwined trimers [19, 20]. The trimer is composed almost entirely of β-sheets arranged in helical barrels, with the distal tip forming a more globular structure that directly contacts the receptor. Consistent with this structure, computational analysis of the secondary structure of J predicts β-sheets across the C-terminal domain, with a single α-helix ∼65 residues from the C-terminus. In T4, the tail fiber turns back on itself to form a six-sheeted, rather than three-sheeted, barrel, suggesting that the α-helix may be part of the distal tip of J.

We decided to interrogate λ host range by generating a library of thousands of genetic variants in the C-terminal domain of J. We first imposed selection for infectivity on wild type (wt) *E. coli*, comparing the patterns of infectivity with the evolution and possible structure of the J protein. We then imposed selection on the same library for infectivity on three λ-resistant hosts, uncovering shared and unique paths to adaptation. By comparing variants within and between conditions, we consider the likely routes by which λ may overcome host resistance, and we ascribe distinct roles for variants that increase promiscuity from those that drive changes to specificity.

## Methods

### Generation of λ variants by codon replacement

We generated a library consisting of most possible single amino acid substitutions in the C-terminal 150 codons of J. We first cloned the wild type J gene onto a high-copy plasmid and used site-directed mutagenesis to disrupt a single BbvCI site in the coding sequence, leaving two same-orientation BbvCI sites on the backbone. We used the BbvCI nickase-based mutagenesis strategy described by Wrenbeck et al. (2016) to randomly replace codons in the targeted region with ‘NNN,’ using 150 degenerate oligonucleotides that each bear homology flanking a single codon [21]. We then inserted a 15 base pair barcode downstream of J and generated a barcode-sequence map by paired-end Illumina sequencing of the mutagenized region, with the barcode on a separate indexing read. We ignored reads with no barcode inserted, barcodes with fewer than five high-quality reads, and barcodes linked to sequences for which more than one read disagreed with the consensus at a given base pair. The barcoded and mapped sequence variants were then cut out of the plasmid backbone using PasI and ligated into PasI-digested and dephosphorylated λ DNA (λ_Sam7/cI857_, NEB #N3011L, Ipswitch, MA, USA). This strain background is obligately lytic at 37°C and has an amber-suppressible lysis system that can infect and kill *E. coli*, but not release progeny, unless the strain is an amber suppressor [22]. The ligated DNA was packaged into virions using the MaxPlax λ packaging extract (Lucigen #MP5120, Middleton, WI, USA), and the virions were sequenced to determine input barcode frequency.

### Selection for infectivity

For each selection, ∼10^7^ virions we mixed with ∼2*10^8^ cells in rich media containing 10 mM MgSO_4_ in 1 mL total volume and incubated at 37°C with shaking at 600 rpm. This binding step was 10 minutes for the wild type host (DH10B) [23] or 60 minutes for hosts with exogenously expressed receptors (DH10B-lamB^Δ^ + p44K-lamB-var). Following binding, the cells were spun down (12,700 rpm × 30 s) and resuspended in LB, and this washing step was repeated twice. The cells were then incubated at 37°C with shaking at 600rpm for an additional 90 minutes to allow infected cells to produce λ progeny, which remained trapped in the cell because DH10B does suppress the Sam7 allele in λ. Cells were spun down, resuspended in Tris-saline-EDTA buffer, and lysed with a combination of lysozyme and bead beating. The lysate was diluted into Tris-saline-magnesium buffer, cleared by centrifugation, and passed through a 0.2 mm filter to remove un-lysed cells. This procedure was repeated for four rounds on the wild type host, or three rounds on hosts with exogenously expressed receptors. At each timepoint, including t=0, the population size was measured by plating on LE392MP cells, which suppress the Sam7 allele, and barcodes were amplified from the population by qPCR for sequencing. Each selection was performed in triplicate.

### Sequencing and scoring of variants

At each timepoint, we deeply sequenced barcodes amplified from the phage populations. We counted only barcodes that were in the barcode–sequence map and had an input read count of ≥20 reads. For each barcode, we estimated the number of progeny produced by the average virion containing that barcode in an infectious cycle (hereafter, progeny). We developed a novel scoring approach, called Model-Bounded Scoring, that directly estimates growth rate, *r*_*var*_, as opposed to commonly used methods that calculate a wild type-normalized enrichment score [24]. By this approach, we measure the population size of the library at each timepoint, and we generate a null model of the expected sequencing counts for each barcode given its starting frequency and assuming no growth. We then estimate the subpopulation size using the sum of the measured and modelled counts and calculate a growth rate by regressing the log subpopulation size over time. We calculate progeny as the exponential of the growth rate, 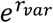, with one progeny meaning the parent survived but did not produce offspring. After four infectious cycles, variant scores were highly correlated between replicates (r^2^ = 0.996, Fig. S1). In this dataset, Model-Bounded Scoring better segregates synonymous and nonsense variants, provides better agreement between replicates, and provides better agreement between different barcodes linked to the same protein sequence than enrichment-based scoring (Fig. S2).

## Results

### Mutational scanning strategy yields infectivity measurements for phages with thousands of J variants

We performed a ‘deep mutational scan’ of phage infectivity in λ by generating a library of phage variants with single amino acid changes in the tail fiber and imposing selection for infectivity on that library (Fig. 2a) [25]. The library consisted of nearly all single amino acid substitutions and several thousand double amino acid substitutions across the C-terminal 150 amino acids of J (Fig. 2b, c). For readability, we numbered the positions starting at the first residue of the mutagenized region. We imposed selection on this library by mixing the phage variants with wild type *E. coli* and isolating intracellular phages after a single round of growth. This selection was repeated for four infectious cycles, and we estimated the abundance of each barcode in the expanding population over time using a novel approach we call ‘Model-Bounded Scoring’ that directly estimates the progeny produced per generation. Barcode abundances between replicates showed strong agreement, as did different barcodes representing the same variant (Fig. 2d, Fig. S2). We used the distribution of nonsense variants and synonymous variants to categorize variant effects. We considered variants with growth rate within two standard deviations of the mean nonsense variant to be ‘null-like’ and those within two standard deviations of the mean synonymous variant to be ‘wt-like’ (Fig. 2e). Variants falling between these two distributions were considered ‘deleterious,’ while variants above the synonymous distribution were considered ‘hyper-infective.’

**Figure 2:**
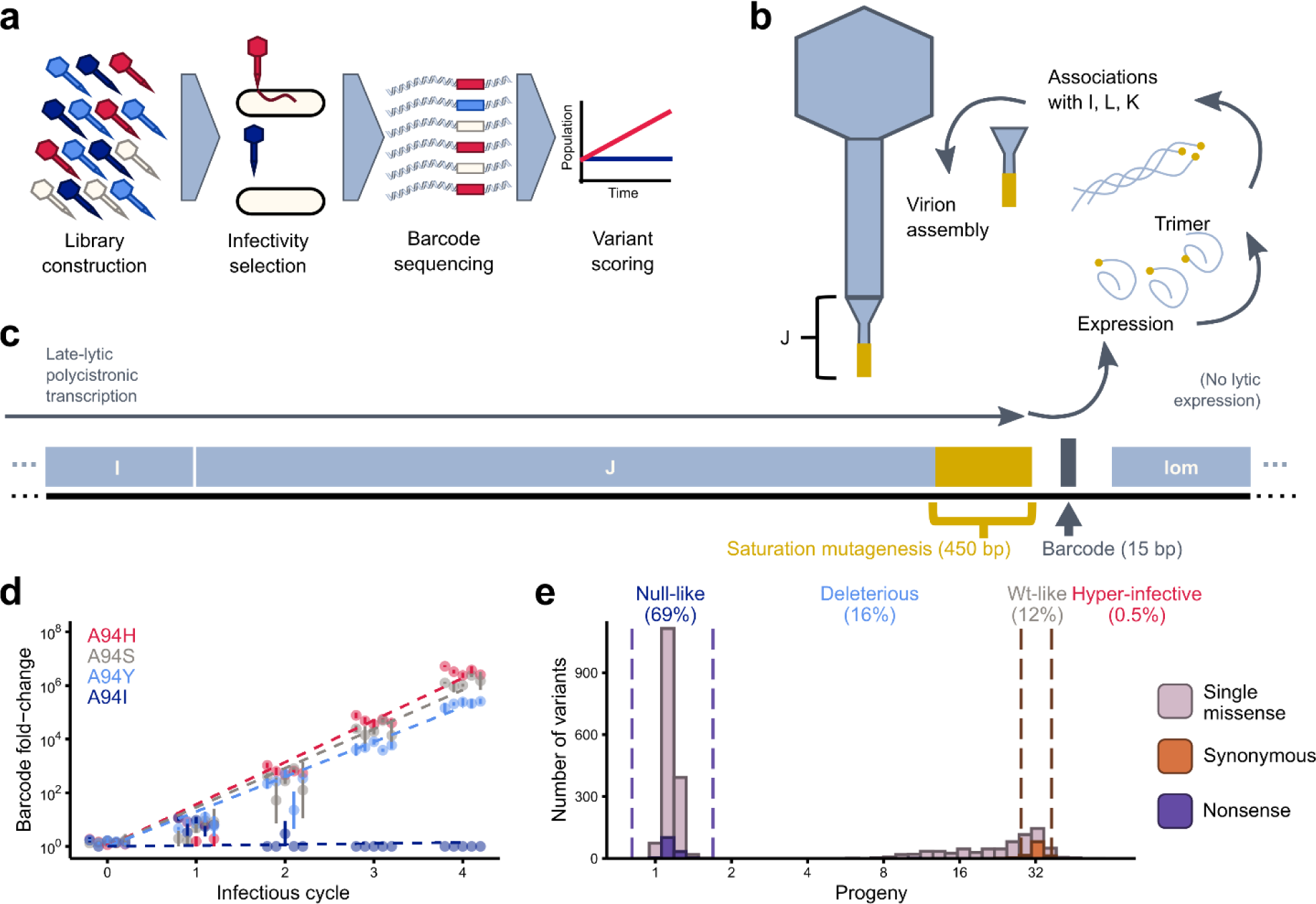
A sequencing-based method to assess phage infectivity across thousands of variants. **(a)** The experimental strategy. A library of phage variants was constructed and subjected to selection for the ability to infect *E. coli*. After each infectious cycle, barcodes corresponding to each variant were deeply sequenced, and their frequencies were used to score each variant using Model-Bounded Scoring. **(b)** Binding of λ to the host is mediated by the J protein, which contacts the receptor, LamB. J spontaneously forms a homotrimer and then associates with accessory proteins that fold J into a mature conformation [47]. **(c)** J is the 3’-most gene on a long polycistronic transcript expressed in late lytic phase that contains most of the capsid genes. We mutagenized the 3’-most 450 bp of J (excluding the stop codon) using NNN codon replacement and inserted a 15 bp barcode downstream of the gene. **(d)** For each of four missense variants at A94, five randomly selected barcodes are plotted by their abundance in the phage population after each infectious cycle relative to the pre-packaging DNA pool. Infectious cycle = 0 corresponds to the packaged but not yet selected phage population. Error bars represent the standard error between 3 replicate selections. For each variant, the growth rate, r, is the slope of an ordinary least squares regression line calculated separately for each barcode and averaged across all barcodes representing the same protein-level variant. The average growth rates for the selected variants are shown by slopes of the dashed lines. **(e)** Progeny per infectious cycle, equal to the exponential of the growth rate (e^r^), is shown for λ bearing synonymous (to wild type) variants in orange, nonsense variants in purple, and single missense variants in mauve. Dashed lines indicate score boundaries used to categorize variants as null-like, deleterious, wt-like, or hyper-infective.

A large fraction (69%) of single missense variants conferred a null-like phenotype (Fig. 3a). Although phage tails rapidly diversify to expand their host range, the fitness landscape of J with respect to its wild type host is much more restrictive than has been seen for other, much more conserved, proteins [26, 27, 28, 29]. The pattern of mutational tolerance in J is broadly consistent with a T4-like barrel structure: extreme intolerance to substitutions to glycine and proline and three regions of high periodicity in mutational tolerance, both of which suggest β-sheet richness (Fig. 3a, S3). We can use this pattern of mutational tolerance to construct a coarse model of J’s structure (Fig. S4), with the receptor-binding portion of the protein encompassing positions ∼50-100. This region of the protein is not, however, enriched for hyper-infective variants. We identified 12 such hyper-infective variants, two of which are accessible through point mutations. That these hyper-infective mutations are found throughout the mutagenized region suggests that residues that do not directly contact the receptor can nevertheless strongly influence receptor binding.

**Figure 3:**
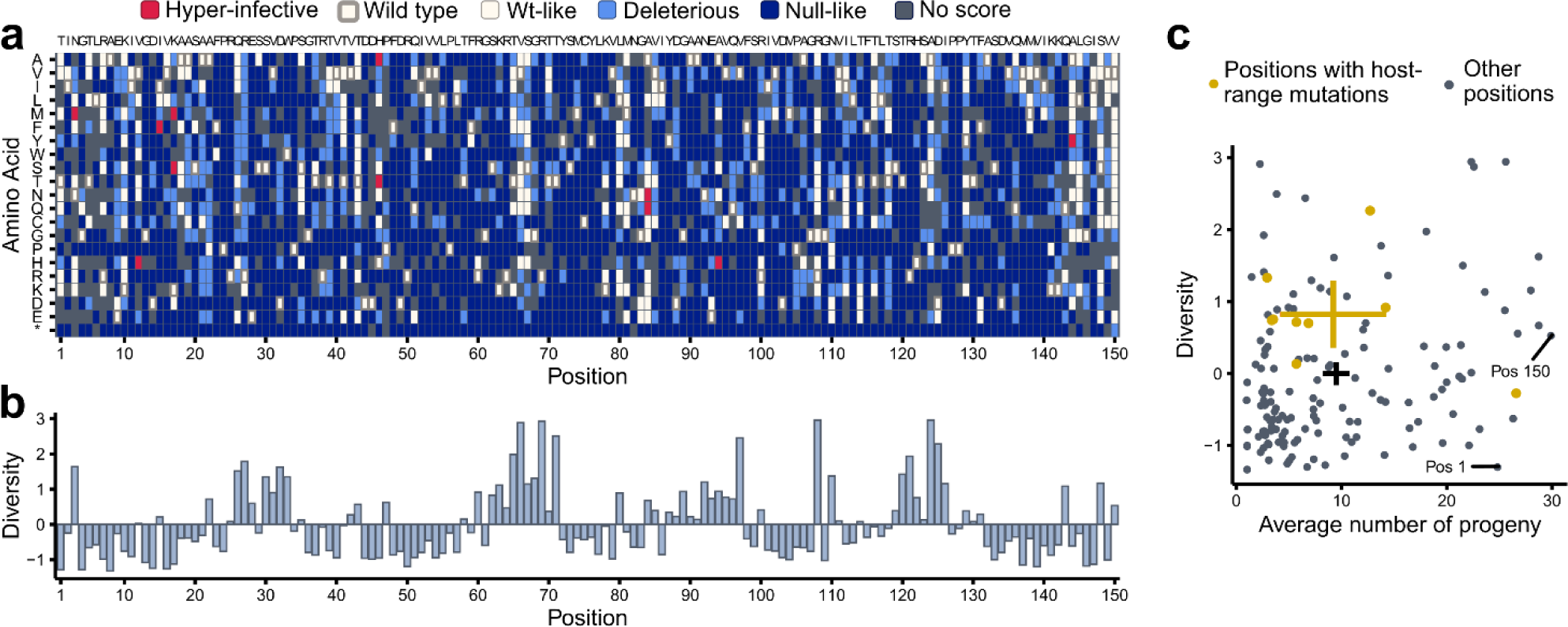
Comparison of empirical mutational tolerance and evolutionary diversity. **(a)** Categorical effects of all amino acid substitutions. Categories are defined in Fig. 2e. The wt sequence of the J region is shown across the top and the amino acid substitutions on the Y axis. **(b)** Amino acid diversity, calculated from ConSurf [48], across 910 orthologs of J. Zero represents the mean diversity across positions. **(c)** For each amino acid position, the evolutionary diversity (y-axis) is compared to the average progeny produced by amino acid substitutions at that position (x-axis). Diversity of each position correlates only weakly with the average number of progeny (r^2^ = 0.06, p<0.01), in contrast to cellular proteins for which mutational tolerance and evolutionary diversity are more strongly related [26, 27, 30]. Gold points indicate positions where mutations have been reported that expand host range [3, 14]; these positions are more diverse but no more mutationally tolerant than the average position. Crosses represent 95% confidence intervals of the mean diversity and progeny of variants for positions with host range mutations (gold) or all positions (black). ‘Pos’: position.

### J fitness landscape is more restrictive than predicted from evolutionary diversity

The extreme intolerance to mutation of J relative to other proteins conflicts with the observation that phage tail fibers, including J homologs, rapidly diversify over evolutionary time. We therefore decided to directly compare the patterns of mutational tolerance from our assay against patterns of mutational tolerance from an alignment of 910 orthologs of J. Positions that harbored hyper-infective variants do not correspond to positions of high amino acid diversity across J orthologs (Fig. 3b). More broadly, the average progeny of variants at a position, a proxy of mutational tolerance, correlated only weakly (r^2^ = 0.06) with the evolutionary diversity of those positions (Fig. 3c) [3, 14]. This observation contrasts with deep mutational scans of conserved cellular proteins, for which much stronger associations are observed between diversity and mutational tolerance [26, 27, 30]. Additionally, the distribution of variant effects argues that J is not trapped on a fitness peak with respect to rapidly binding to its wild type host. Point mutations were slightly less likely to be strongly deleterious than all possible amino acid substitutions, even when synonymous mutations are excluded (Fig. S5). Moreover, the best-scoring point mutation, I15F, was 4.2 standard deviations above the distribution of synonymous variants, producing 35% more progeny per generation than the wild type. Thus, the selection pressures that have shaped the evolutionary history and the wild type sequence of λ may have acted on a different property than is being selected for in our assays.

In J, peaks of diversity (Fig. 3b) that corresponded to many wild type-like variants (*e*.*g*. positions 67 or 108) can be separated from those that corresponded to many deleterious or null-like variants (*e*.*g*. positions 30 and 97). Furthermore, λ host range mutations tended to fall at these diverse, mutationally intolerant positions (gold points, Fig. 3c). We posit that by imposing selection for infectivity on a single host, we measured a selective pressure that is too narrow to adequately reflect λ’s entire evolutionary history on multiple hosts. Positions important for mediating infection of a single host would be mutationally intolerant in our assay, as the amino acid optimal for binding wild type LamB would likely have already been fixed in λ. However, these same positions could be under diversifying selection over evolutionary time during which the host has varied. Therefore, a complete understanding of λ evolution requires comparing the effects of J variants across many hosts.

### Adaptation to a set of resistant hosts

To investigate mechanisms of adaptation to a resistant host, we challenged the library of J variants with a set of *E. coli* hosts bearing novel λ resistance mutations. A deep mutational scan of LamB, the λ receptor, showed that many point mutations specific for λ resistance are in or proximal to loop L6, the presumptive λ binding site [31]. We chose three receptor mutations, LamB-R219H, LamB-T264I, and LamB-G267D, that confer λ resistance but allow maltodextrin transport, on the basis that these mutations specifically disrupt the λ binding site rather than disrupting stability of the receptor. In a *lamB*-deletion background, we generated a *lac*-inducible expression vector for the wild type and the three *lamB* alleles, which were expressed using 0.1 mM IPTG. We imposed selection for infectivity on the λ library on each of the four *E. coli* strains, and we estimated progeny using Model-Bounded Scoring.

Similar to selection on wild type *E. coli*, synonymous J variants were more infective than nonsense variants on each of the three resistant *E. coli* strains (Fig. 4a). Because Model-Bounded Scoring estimates real growth rate, rather than relative fitness, we can directly compare the growth of λ bearing the same J variants on different hosts without normalizing to a reference allele. λ was slightly more infective on the IPTG-expressed wild type receptor than when this receptor was expressed from the endogenous *lamB* locus (31 progeny vs. 27 progeny for wild type J), but infectivity of the variants was highly correlated between the selections (r^2^ = 0.958, Fig. 4c). The resistant *lamB* alleles did not confer absolute resistance, but decreased the progeny produced by wild type λ from 31 (LamB-wt) to 22 (LamB-G267D), 2.7 (LamB-T264I), and 2.0 (LamB-R219H). For all three *lamB* mutants, we could identify single missense variants in J that restored progeny to 40-100% of what was produced on the wild type host (Fig. 4a). Moreover, a much larger fraction of variants significantly outperformed the synonymous distribution on each resistant host. Adaptive variants in each case were broadly distributed over the sequence, not highlighting a domain or structural feature uniquely necessary for adaptation (Fig. 4b). With some exceptions, adaptive variants were neutral-to-beneficial at infecting LamB-wt (Fig. 4c). However, the reverse was not true; variants that were highly infective on the wild type receptor were frequently less infective on the novel hosts.

**Figure 4:**
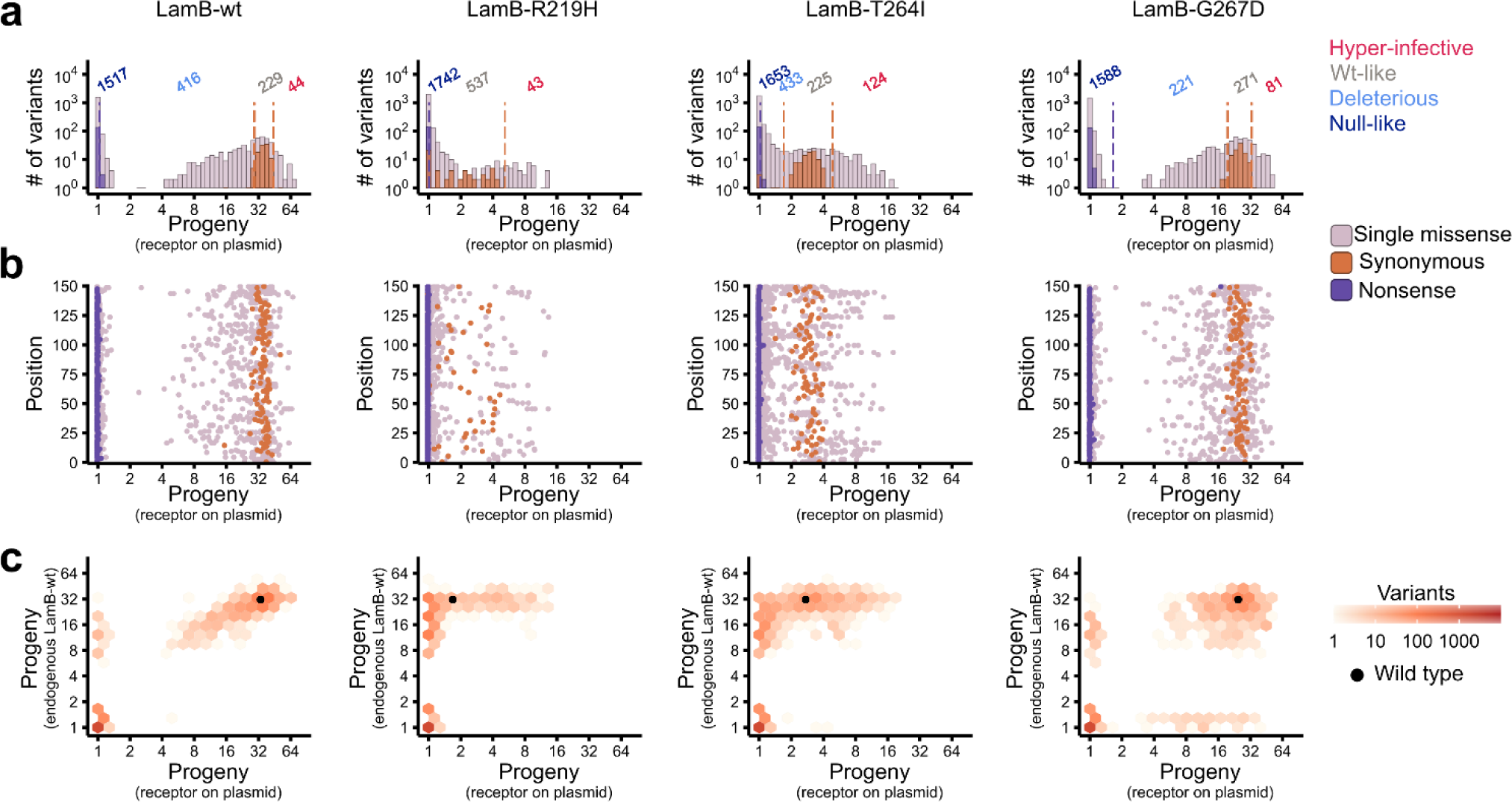
Selection of λ bearing J variants on λ-resistant hosts. **(a)** Distribution of fitness effects on each host. In each case, the synonymous and nonsense distributions can be separated, despite the nominal λ-resistance of each lamB allele. Variants that outperform wild type λ do so by a larger margin on hosts that are more resistant. **(b)** Progeny for each J variant shown by the position of the mutation in the sequence. Adaptive mutations occur frequently at a subset of positions, but these positions are spread over the entire mutagenized region. Positions with many hyper-infective mutations are more apparent on the more resistant hosts LamB-R219H and LamB-T264I than on LamB-wt or LamB-G267D. **(c)** Correlation between progeny produced by λ bearing each J variant on its wild type host (y-axis), or on a host bearing a plasmid-borne lamB allele (x-axis). Variants that are highly infective on a non-wt host tend to also be infective on the wild type host, with some exceptions. However, many variants that are highly infective on the wild type host are poorly infective on non-wt hosts.

### Specific and general mechanisms of adaptation

A comparison of the infectivity of J variants between two hosts shows that the most infective J variant with respect to one host was frequently highly infective on the other host as well (Fig. 4c, S6). In cases for which a J variant conferred infectivity on one host but not the other, it nearly always conferred infectivity on the host that is less resistant to wild type λ. Reciprocally, for hosts with greater resistance to wild type λ, a smaller fraction of J variants conferred infectivity (*i*.*e*., were not null-like). This patten is consistent with a ‘nested’ model in which each host and pathogen has a set amount of resistance or counter-resistance that is not dependent on the other player (Fig. 5a). In this model, infectivity is determined by the relative strength of resistance and counter-resistance. This model would contrast with a ‘lock-key’ model [34], in which infectivity is determined by how a well a given λ variant matches a given receptor (Fig. 5b). The relevant distinction is that under a nested model, but not a lock-key model, a J variant may gain generic counter-resistance that improves its infectivity on many hosts.

**Figure 5:**
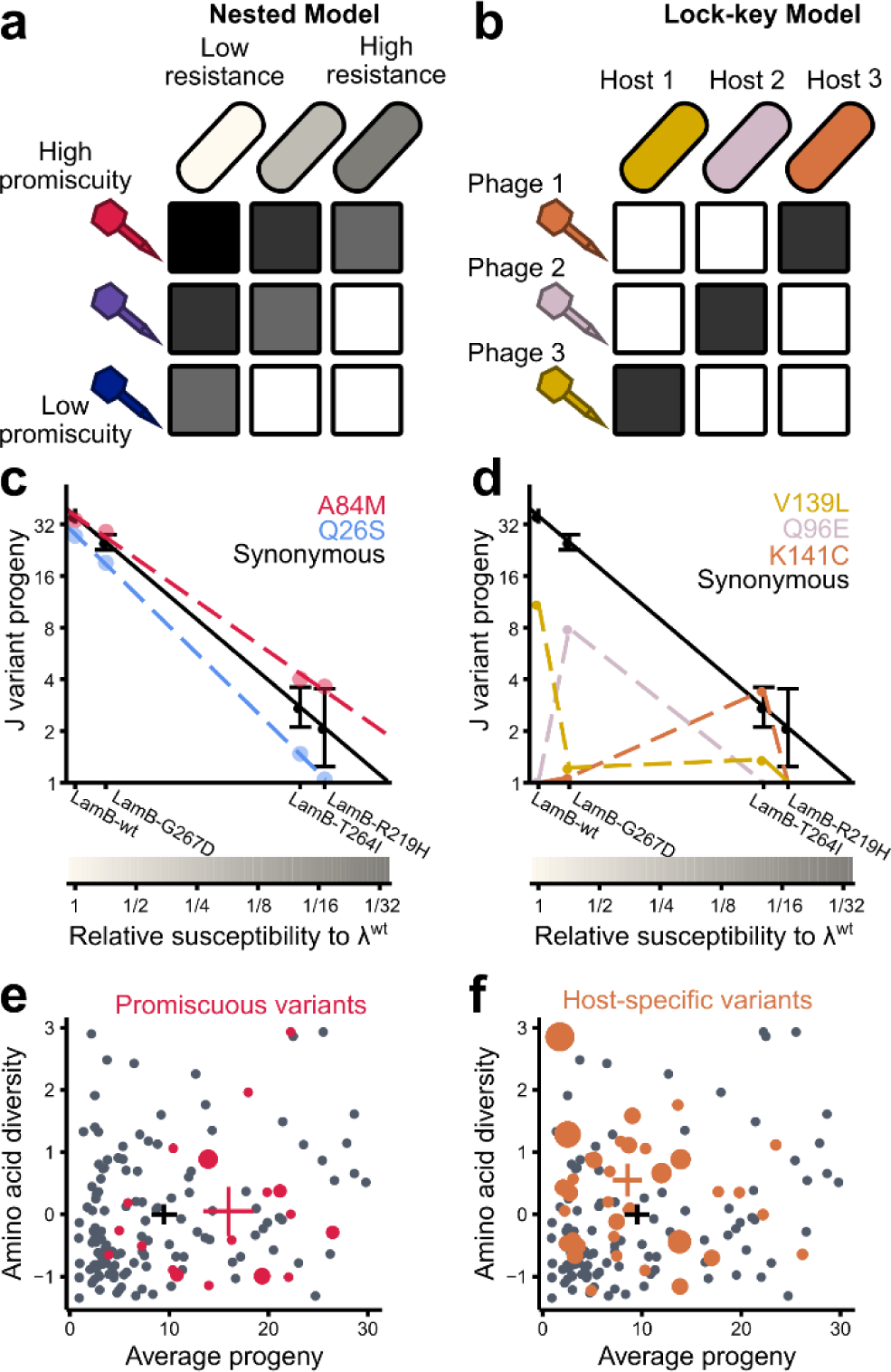
Discrimination of promiscuity from specificity by comparison of variant effects across different hosts. **(a)** Under a nested model, phages have more or less ability to generally infect hosts, which we describe as ‘promiscuity,’ which contends with the level of resistance of potential hosts. **(b)** Under a lock-key model, the specific relationship between a host and phage determines infectivity, rather than host-independent properties of the phage or phage-independent properties of the host. **(c)** We can estimate the level of resistance of a given host by asking how well λ^wt^ produces progeny on it compared to LamB-wt (x-axis). Some J variants, like A84M are less affected by resistance than wild type λ, whereas others, like Q26S, are more affected. We calculate promiscuity as the area under the curve for a regression line comparing the variant infectivity to host susceptibility, relative to wild type λ. The black line represents synonymous variants, with error bars equal to +/- 1 standard deviation. **(d)** Some J variants confer infectivity on only a single host and are null-like on all others. Most of these variants, like Q96E, are deleterious, even on the host for which they are specific. Therefore, most of these variants cannot drive adaptation to a novel host by themselves, though they may work in concert with other variants. **(e)** Positions with promiscuous variants are shown in red with respect to their tolerance to mutation and amino acid diversity, with the size of the circle representing the number of unique promiscuous variants. The weighted average of these positions (cross, red) is more tolerant to mutation but not more diverse than the average of all positions (cross, black). Crosses represent 95% confidence intervals. **(f)** Variants that display specific infectivity on a single LamB variant fall at positions shown in orange. The weighted average of these positions has higher amino acid diversity, but is not more mutationally tolerant, than the average position.

The property of generic counter-resistance can be compared to the property of enzyme ‘promiscuity,’ in which enzymes weakly catalyze non-canonical reactions in addition to their normal biological activities [35]. Under the right selection pressure, mutations can expand or improve these promiscuous activities, and evolution may eventually lead to specialization in the new activities [35]. By analogy, J might be said to have a baseline promiscuity in that it can weakly bind non-LamB-wt receptors, and that variants have increased promiscuity if they improve infectivity broadly over the space of potential hosts, rather than only on LamB-wt (Fig. 5c). To estimate the promiscuity of J variants, we plotted in log-space the progeny conferred by a variant on each host, compared to the susceptibility of these hosts to wild type λ. For synonymous variants (Fig. 5c, black line), this relationship is 1:1 by definition. However, some variants, like A84M, have a shallower slope indicating that they were less affected by the resistance of receptor mutations; we call variants with significantly greater area under the curve than synonymous variants ‘promiscuous.’ Most promiscuous variants grew well on the wild type host (Fig. S8), implying that these promiscuous variants could persist in a λ population prior to encountering a resistant host. However, that these variants have not already been fixed in the population may imply subtle costs to promiscuity, such as thermodynamic instability [35, 36].

Additionally, 49 single missense variants conferred growth on only a single host, most of which conferred growth on LamB-G267D (such as Q96E in Fig. 5d, S9). With a few exceptions, like K141C (Fig. 5d), these host-specific variants did not confer greater infectivity than wild type J on any host we tested; they were merely dramatically more infective on a particular host compared to the other hosts. Thus, while these host-specific variants may be an important part of adapting to a host, they are generally insufficient to directly overcome resistance. Rather, we posit that the acquisition of host-specific variants may follow after the acquisition of promiscuous variants and either ameliorate thermodynamic costs or prevent off-target binding. This process could explain why very few LamB-wt-specific variants were observed, and why they were less infective than variants specific to other hosts (Fig. S10), as if the wild type J sequence has already been selected for near-maximal specificity to its host.

Based on our analysis of J variants growing on a wild type host, positions with low mutational tolerance but high diversity over evolutionary time are predicted to be enriched within variants that drive adaptation to a novel receptor (Fig. 3c). Contrary to expectations, promiscuous variants tended to occur at positions with higher-than-average progeny across variants (Fig. 5e). By contrast, host-specific variants tended to occur at positions that are intolerant to mutation and diversified over evolutionary time (Fig. 5f). Thus, while our hypothesis that these mutationally intolerant, evolutionarily diverse positions are driving host-specificity is largely supported, host-specificity is not equivalent to overcoming resistance, which can happen through host-nonspecific mechanisms (*i*.*e*., promiscuity). This distinction implies that host range mutations arising in experimental evolution studies have not generally been promiscuous variants, as these host range mutations have mostly fallen at positions both mutationally intolerant and diverse (Fig. 3c). In our analysis, many of these host range mutations were null-like on all the hosts tested, including LamB-wt (Table S1), suggesting that the effects of these variants may be specific to the hosts used in those studies and/or co-occurring mutations in J.

### Positive epistasis potentiates adaptation to a new host

In addition to single missense variants, the J library contained approximately 7,500 variants with two missense mutations, sparsely surveying the space of >4 million possible double missense variants. We wondered whether combinations of promiscuous and host-specific mutations could help mediate adaptation to a novel receptor beyond what either class of mutations would confer in isolation. For example, A94S is a promiscuous variant, but was mildly impaired for infectivity on LamB-G267D, and S30W is a LamB-G267D-specific variant that was mildly impaired on LamB-G267D but null-like on all others (Fig. 6a). However, the double missense variant A94S+S30W had both high growth and moderate specificity on LamB-G267D. This double missense variant therefore exhibits positive epistasis on one host, though it is poorly infective on the other hosts.

**Figure 6:**
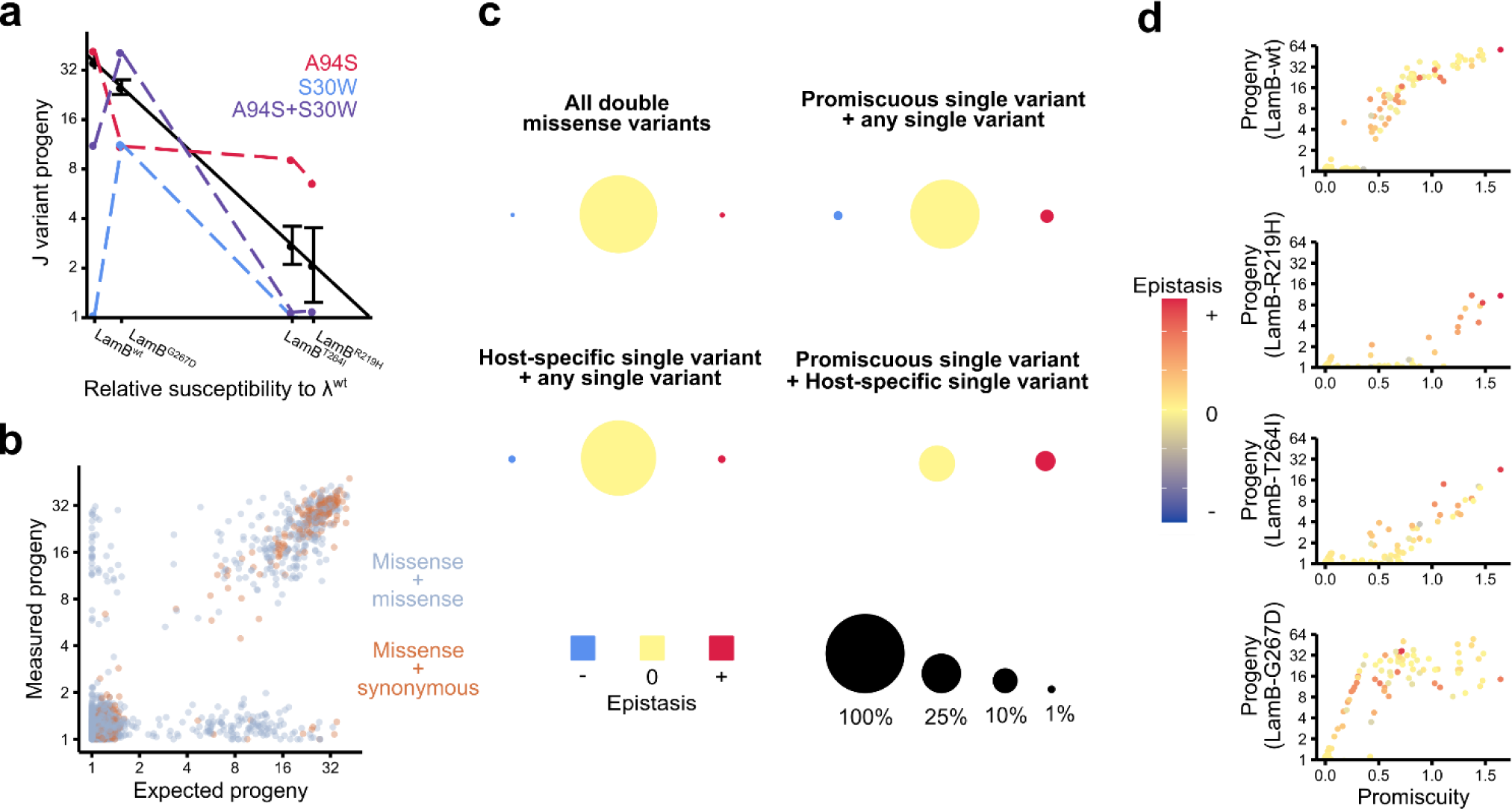
Double missense variants in J can mediate adaptation to new hosts. **(a)** The double missense variant S30W, A94S is a combination of a promiscuous variant (A94S) and a LamB-G267D-specific variant (S30W). The double missense variant exhibits sign epistasis, improving infectivity on LamB-G267D despite the deleteriousness of each single missense variant. **(b)** For double missense variants (silver), we calculated the expected progeny from the progeny of each of the single missense variants using a simple multiplicative model. A subset of double missense variants strongly deviates from the multiplicative model, in contrast to variants in which a single missense mutation is paired with a single synonymous mutation (orange), which are more likely to agree with the multiplicative model (r^2^ = 0.92 missense x synonymous vs. r^2^ = 0.64 missense x missense). **(c)** Across the four hosts, most double missense variants in J do not exhibit significant epistasis (top left panel). However, double missense variants that contain a promiscuous variant, a host-specific variant, or both, are more likely to exhibit significant epistasis. We measured significant epistasis in 13.2% of double missense variants containing promiscuous variants, 5.4% containing host-specific variants, and 23.8% containing both, compared to 1.8% containing neither. **(d)** For each host, progeny is positively associated with promiscuity and with positive epistasis. However, these associations become more salient on resistant hosts, with all infective variants on LamB-R219H being promiscuous and positively epistatic, compared to a minority of variants on LamB-wt.

On LamB-wt, only 6.2% of double missense variants were infective (*i*.*e*., not null-like), compared to 26% of single missense variants that were infective. For each double missense variant, we calculated the expected progeny given the progeny from each single variant, using a simple multiplicative model. Most variants yielded similar values for measured and expected progeny, but ∼45% of infective variants (2.7% of total variants) exhibited significant epistasis (Fig. 6b), including 22 infective variants for which both mutations were null-like on their own. These variants with dramatic reciprocal sign epistasis were enriched for positions with host range mutations [3, 14], which appeared 20 times among these 22 variants. As a control, we also considered variants in which a single missense mutation was paired with a synonymous mutation. Such variants exhibited better agreement between expected and empirical scores (r^2^ = 0.92) than double missense variants (r^2^ = 0.64), suggesting that the prevalence of epistasis in double missense variants is not an artifact of the fact that most double missense variants were less abundant in the library than single missense variants.

Similar to the selection on the wild type host, strong positive epistasis was prevalent on each resistant host. Single missense variants that conferred promiscuity frequently interacted epistatically with other variants (Fig. 6c). Of double missense variants that contained a promiscuous single missense variant, 73/550 (13.3%) exhibited significant epistasis compared to only 2.2% of all double missense variants. The rare cases of double missense variants consisting of one promiscuous variant and one host-specific variant were even more likely to exhibit epistasis (5/21, 23.8%). In these five examples, epistasis was always in the positive direction.

Additionally, the effects of positive epistasis became more salient when λ was challenged with a resistant host. Double missense variants that were infective on LamB-wt had varying levels of promiscuity and positive epistasis (Fig. 6d). By contrast, double missense variants that were infective on the most resistant host, LamB-R219H, nearly exclusively had both high promiscuity and positive epistasis. The other two hosts revealed intermediate effects.

## Discussion

Although phages and bacteria are often presumed to coevolve stably and indefinitely, an “asymmetry in coevolutionary potential of these hosts and parasites” exists [5]. To investigate how λ overcomes host resistance, we analyzed thousands of variants of its tail fiber protein, J, on a small set of resistant *E. coli* hosts. We find that promiscuous J variants, which increase infectivity on a broad range of hosts, underlie the re-establishment of infection on resistant hosts. These variants co-exist with other, host-specific, variants that generally do not increase infectivity on any host but have smaller losses to infectivity on a single host. We posit that both types of variants are important for a phage to adapt to a new host, with host-specific variants likely ameliorating costs associated with promiscuity. This framing has implications for experimental evolution studies, protein adaptation more broadly, and natural phage-bacteria communities.

When phages and bacteria cyclically develop resistance and counter-resistance in experimental evolution studies, this coevolution is frequently characterized by an initial escalation of both host resistance and phage counter-resistance. This escalation eventually reaches an asymptote and is followed by negative frequency-dependent selection (“Kill the winner” dynamics) in which the dominant phages are most infective on the most common hosts [6, 7]. This pattern is well explained by a model in which promiscuous variants drive broadened host range but come at a cumulative cost, manifesting as lower growth rate relative to their host-specific counterparts [37]. The sequential acquisition of promiscuous variants could also open up pathways for a phage to infect a highly resistant host by first adapting to a less resistant host in the same environment. For example, Werts *et al*. (1994) could not directly isolate λ that overcame the resistance allele LamB-G151D, but by pre-adapting the phage to other hosts with weaker resistance, they found double mutants able to grow on LamB-G151D [14]. However, thermodynamic costs associated with promiscuity may also constrain paths to adaptation. In their isolation of LamB-independent λ strains that use OmpF as receptor, Meyer *et al*. (2012) repeatedly identified the same set of 4-5 adaptive variants across dozens of independent cultures [3], suggesting considerable constraint on the path to counter-resistance [38]. In a follow-up study, the bi-specific intermediate, which binds both LamB and OmpF, was less stable than the LamB-specific parent, and selection for OmpF specificity was sufficient to restore stability [39].

In the context of enzymes, promiscuous activities provide convenient starting points for adaptation to novel protein functions [35]. Increasing the stability of the parent enzyme can potentiate greater adaptation by compensating for the mild destabilization associated with some adaptive variants [36, 40]. Similarly, the low level of infectivity mediated by wild type J on resistant hosts serves as a starting point for adaptation to those hosts. Second mutations can compensate for the initial mutations that confer promiscuity, offsetting the potential costs of these initial mutations and resulting in highly infective and/or promiscuous double variants. This effect can be seen in the strong association between positive epistasis and promiscuity, whereby promiscuous single variants were more likely to have positive epistatic interactions than non-promiscuous variants (Fig. 6c) and rare promiscuous variants that drive adaptation to LamB-R219H were always positively epistatic (Fig. 6d). At a mechanistic level, promiscuous J variants may shift between multiple semi-stable protein conformations, or they may heterogeneously fold into one of multiple stable conformations. The latter mechanism was found to underlie the LamB-OmpF bi-specific intermediates characterized by Petrie *et al*. (2018) [39]. Under a model requiring multiple protein conformations, destabilization may be fundamentally linked to promiscuity rather than incidental to it. Therefore, counter to observations with promiscuous enzymes [36, 40], stabilizing mutations in phages are unlikely to precede adaptive ones. Instead, a destabilizing mutation that increases promiscuity must come first, followed by a compensatory mutation that re-stabilizes the protein into an optimal conformation for infection of the most abundant host. This prediction would also explain why J is so broadly intolerant of mutation: repeated evolutionary transitions between stable and unstable sequences leave J close to a threshold of severe destabilization, compared to proteins whose evolutionary histories are dominated by stable sequences [41, 42].

Naturally occurring phage–bacteria interactions also show patterns consistent with a balance between host-specific and promiscuous variation. Phages and bacteria isolated from the same environment form infection networks exhibiting both nestedness and ‘modularity,’ a property of lock-key models, with nestedness dominating at small scales involving highly related strains [43, 44]. This pattern is consistent with promiscuity driving counter-resistance to a newly resistant host, and modularity arising between more diverged hosts. The extent to which the evolution of phages involves adaptation to diverged hosts remains unclear. Orthologs of both J and LamB are broadly distributed among enterobacteria, and even appear in distant ε-proteobacteria species, suggesting either an ancient origin of this host–pathogen relationship or frequent cross-taxa jumps in host range. However, the limited breadth of hosts in which nestedness is observed suggests practical constraints to the promiscuity of a phage tail: a single promiscuous phage variant is more likely to be infective multiple receptor variants within a single host species than on receptors of multiple related host species. In our hands, an *E. coli* host expressing a LamB ortholog from another enterobacteria (*C. freundii, Y. pestis*, or *S. marcescens*) did not support growth of our library, which mostly contained single missense mutations, suggesting that multiple mutations may be required for jumps between species.

We conclude that although λ faces significant evolutionary hurdles not faced by its host, it can establish common paths to adaptation on multiple potential hosts by maintaining a balance between promiscuity and host-specificity. This balance may be mediated by mild destabilization of the protein, allowing it to sample multiple conformations, although further work is needed to directly test this hypothesis. Although we surveyed adaptation to only a small set of resistant hosts, this general framework is consistent with prior observations of how λ evolves to switch to a novel receptor. This framework may apply broadly to other phages and viruses for which mutations are more difficult to assay *en masse* [7, 45, 46].

## Acknowledgements

We thank Benjamin Kerr for his advice on the manuscript, and Alan Rubin and Matt Rich for scripts that facilitated filtering and counting barcode sequences. This work was supported by grants T32 HG000035 and RM1 HG010461 from the National Human Genome Research Institute of the NIH.

## Ethics Declarations

Conflicts of Interest: The authors declare that they have no conflicts of interest.

