## Supplementary material for "Balance between promiscuity and specificity in phage λ host range"

### Supplementary Methods

#### Library generation

The library of λ variants was generated by first digesting the λ genome with PasI, then cloning the small fragment that contains all of J plus some flanking DNA, onto the high-copy *E. coli* vector p44K. This vector contains three BbvCI sites, with two in the forward and one in the reverse orientation. Since all BbvCI sites must be in the same orientation for Nicking Mutagenesis, the reverse site was altered at two positions using site-directed mutagenesis. This BbvCI site was in the J coding sequence, and both edits were synonymous changes. The entire J coding sequence was subsequently confirmed by Sanger sequencing. We performed Nicking mutagenesis as described by Wrenbeck *et al.* (2016), using 150 primers, each containing a random (NNN) codon. We performed one round of mutagenesis, in three pools of 50 primers each, and Sanger sequenced colonies to check the level of mutagenesis. There was a high carryover of the wild type sequence (~30%), so we pooled the libraries and performed an additional round of mutagenesis. After the second round, we determined that the library contained mostly single NNN replacements, with some double mutations and ~12% wild type. This library was used for all experiments.

To barcode the library, we amplified the $Kan^r$ gene from pET-9a using primers that added BamHI sites at each end, a 15-base barcode at the 5' end, and homology to the vector on both ends. We digested the p44k-borne library with HinDIII and used Gibson assembly [Gibson et al. 2009] to insert the $Kan^r$ amplicon, selecting on 50 mg/mL kanamycin to remove the parent plasmid. The kanamycin cassette was removed by digesting the plasmid with BamHI, cutting the fully cut product out of an agarose gel, and performing a self-ligation with T4 DNA ligase. The ligation product did not produce any $Kan^r$ colonies.

To computationally link barcodes and variants, we amplified from immediately upstream of the mutagenized region to immediately downstream of the barcode and sequenced the amplicon on the Illumina MiSeq platform. Consensus reads of at least five barcodes were constructed, conditional on >80% agreement between reads, and assigned to barcodes. These reads were also used to calculate input frequencies for each barcode for the purpose of determining input read thresholding and expected barcode counts. We measured input frequency (infectious cycle = 0, the phage population prior to selection) separately for each replicate by amplifying barcodes from the input phage population, and these replicate-specific frequencies were used for determining scores for variants.

Prior to each selection, a midiprep of the plasmid borne library was digested with PasI and ligated with PasI-digested de-phosphorylated λ genome. The ligation product was packaged into λ using the MaxPlax λ packaging extract. The variants prior to the first infectious cycle, while containing the correct DNA variants, therefore have a wild type J protein. The wild type J protein is replaced with the variant protein at the first infectious cycle.

#### Strains and plasmids

Selections were performed in the background of DH10B. For the first selection, the DH10B cells were grown from ElectroMax cells (Thermo-Fisher). For the second set of selections, we generated DH10B-lamB$^{\Delta}$, which harbors an in-frame scarless deletion of the lamB gene. Briefly, pSIM5 was transformed into DH10B and the transformed strain was heat shocked at 42$^{O}$C to induce expression of the λ-red recombination system, then made electrocompetent. A linear cassette containing tetA and sacB flanked by homology to the lamB locus was then transformed in, and cells were selected on tetracycline. pSIM5, which was lost during heat shock, was retransformed, and the cells were heat shocked and made electrocompetent. Finally, an oligo flanking both sides of the lamB locus was transformed in and cells were selected on sucrose + fusaric acid, which selects against the tetA-sacB cassette. Clean deletion of the lamB gene was confirmed by Sanger sequencing and pSIM5 was cured by growing cells at 37$^{O}$C overnight and choosing a chloramphenicol-sensitive colony by replicate streaking on LB and LB+chloramphenicol.

For the second set of selections, we expressed lamB variants from the high-copy expression vector p44k. We used PCR to amplify lamB from the DH10B genome, and cloned it into p44K such that the gene is *lac*-inducible, producing p44K-lamB. We performed site-directed mutagenesis to separately introduce the alleles g659a(R219H), c794t(T264I), and g803a(G267D), and the mutations were confirmed by Sanger sequencing the entire lamB coding sequence. We transformed each lamB variant into DH10B-lamB$^{\Delta}$.

We used λ DNA from NEB, which contains the cI857 and Sam7 alleles. cI857 confers temperature sensitive lysogeny and is obligately lytic at 37$^{O}$C, at which experiments were performed. Sam7 confers amber-suppressible lysis. In DH10B, $\lambda_{Sam7}$ produces intracellular progeny but does not lyse cells following infection. However, progeny can be released if cells are lysed exogenously. We used the amber-suppressor strain LE392MP (Lucigen) for measuring phage stocks.

**Media and expression**

Prior to each selection, we prepared cells for infection. Cells were grown overnight at 37$^{O}$C and 250 rpm in LB + 10 mM $MgSO_4$ + 0.2% maltose for DH10B or DH10B-lamB$^{\Delta}$, or the same media with 100 mg/mL ampicillin and 0.1 mM IPTG for DH10B-lamB$^{\Delta}$(p44K-lamB). Cells were back-diluted 1:100 into the same media in the morning, grown for 4 hours, pelleted and resuspended into 10 mM $MgSO_4$ to a volume such that $OD_{600nm}$ = 0.5. Cells were stored this way at 4$^{O}$C for up to 72 hours, and plated at a $10^{-6}$ dilution on LB immediately prior to selection to determine viability.

**Selection conditions**

For selections, 1 mL of cells were pelleted and resuspended in 1 mL LB + 10mM $MgSO_4$ + 0.2% maltose. As a negative control, we included a tube containing DH10B-lamB$^{\Delta}$, which produce very few (though non-zero) phage progeny. The phage library was added, using >$10^6$ pfu (plaque-forming units), and the culture was shaken at 37$^{O}$C and 600 rpm for the prescribed binding time (10 minutes for the first selection or 60 minutes for the second set of selections). Cells were pelleted, washed twice in 1 mL LB, then grown at 37$^{O}$C and 600 rpm for 90 minutes.

Following this growth, cells were manually lysed as follows. Cells were pelleted and resuspended in 100 mL STE (100 mM NaCl + 10 mM Tris + 0.5 mM EDTA). 0.1 µL ReadyLyse lysozyme (Lucigen) was added and cells were incubated for 10 minutes at room temperature. 250 mg of fine glass beads were added and the mixture was shaken in a bead beater for 30 seconds at 4$^{O}$C. Lysate was recovered by adding 500 mL TMS (100 mM NaCl + 10 mM Tris + 10 mM $MgSO_4$), vortexing, letting the beads settle, and pipetting off the supernatant. The lysate was cleared by centrifugation (12,700 rpm x 2 minutes), and the cleared lysate was driven through a 0.2 mm filter to remove any remaining cells.

**Sequencing library preparation**

At each replicate and for each timepoint, 5 mL lysate was used as the template for SYBR-based qPCR using primers that add sequencing adapters and custom indices. Because of the high magnesium content of the lysate, we used Phusion with the 5x GC buffer, and added DMSO to 3% final concentration. The barcodes were amplified and monitored manually, with tubes removed as they entered exponential phase. All tubes were removed by cycle 15. DNA was cleaned up from each reaction on a Zymo clean & concentrator column, the product concentration was determined by Qubit, and libraries were mixed at equimolar ratio and prepared for Illumina NextSeq sequencing.

**Sequencing and barcode counting**

Amplicons containing the indexed barcodes were deeply sequenced by the Illumina NextSeq, yielding at least 10x average coverage of the library for each replicate and timepoint. The sequences were trimmed down to just the barcode, and they were counted using Enrich2 with no filters. The counts files were then used for Model-Bounded Scoring.

**Model-Bounded Scoring**

We developed a custom scoring pipeline, which we call "Model-Bounded Scoring." The name refers to the fact that each barcode is initially modeled as a zero-fitness variant, with the model placing an effective lower bound on the score a variant can receive. For variants that have non-zero counts at later timepoints, the model does not substantially alter the score.

For each barcode, we want to estimate the growth rate, $r_v$. Typically, we would attempt to estimate the number of viruses containing that barcode at each timepoint, $n_{v,t}$, compared to the number of viruses containing that barcode at t=0, $n_{v,0}$. We could then solve for $r_v$ using the equation:

$$e^{r_v * t} = \frac{n_{v,t}}{n_{v,0}} \quad \{1\}$$

Assuming each barcode has a minimum growth rate of zero, $n_{v,t}$ should always be positive, and this equation will hold. However, when $n_{v,t}$ is estimated from counting barcodes, it is common for many barcodes to have zero counts, which will generally cause $n_{v,t}$ to be estimated as zero. Since log(0) cannot be usefully evaluated, this estimate makes it impossible to come up for a useful estimate of $r_v$. A common way to address this problem is to add a constant positive value, termed a 'pseudocount' to each barcode count, therefore causing $n_{v,t}$ to always be positive. However, this can have distortionary effects on the scores.

Instead of trying to solve equation 1 as written, we assign each barcode a modelled growth rate, $r^{\Psi}$. We assign $r^{\Psi} = 0$ for all barcodes, but it is not strictly necessary to do so. We then try to solve the equation:

$$e^{(r_v + r^{\Psi}) * t} = \frac{n_{v,t} + n^{\Psi}_{v,t}}{n_{v,0} + n^{\Psi}_{v,0}} \quad \{2\}$$

Since we assign $r^{\Psi} = 0$, the left half of equation 2 is equivalent to the left half of equation 1. Values for n can be estimated from sequencing counts as follows:

$$n_{v,t} = \frac{c_{v,t} * N_t}{C_t} \tag{3}$$

where $c_{v,t}$ is the counts of the barcode, v, at a given timepoint, t; $C_t$ is the total counts of all barcodes at that timepoint; and $N_t$ is the total population size at that timepoint. For the modelled counts, we estimate as follows:

$$n^{\Psi}_{v,t} = n^{\Psi}_{v,0} e^{r^{\Psi} * t} \tag{4}$$

Combining these equations, we can solve for:

$$r_v * t = ln\left(\frac{c_{v,t} * N_t * C_0}{c_{v,0} * N_0 * C_t} + e^{r^{\Psi} * t}\right) + ln(1/2) \tag{5}$$

In practice, rather than solving this equation separately for each timepoint, we regress the right half of the equation over time and estimate $r_v$ as the slope of the regression. Since ln(1/2) is constant at every timepoint, it does not contribute to the slope. Thus, the term we actually compute is

$$ln\left(\frac{c_{v,t} * N_t * C_0}{c_{v,0} * N_0 * C_t} + 1\right) \tag{6}$$

which is regressed over time with an intercept at the origin.

We calculate slope as described separately for each barcode and each replicate. We also record the standard error of the estimate from the regression, which is propagated to form the error term.

This calculation requires an estimation of the population size, $N_t$, which we measured by plating the library at each timepoint. In some experimental designs, however, library sizes are not estimated. Unlike enrichment scoring, Model-Bounded Scoring does not normalize all scores to a reference allele and attempting to do so is not recommended. If the growth rate of wild type (or another reference allele) is known, the population size can be estimated from the growth rate, and the appropriate term to be regressed over time is:

$$ln\left((2e^{r_{wt} * t} - 1)\frac{c_{wt,0} * c_{v,t}}{c_{wt,t} * c_{v,0}} + 1\right) \tag{7}$$

In our dataset, using this approach slightly improved replicability between different barcodes, but substantially decreased the linearity of each barcode over time (the improvement in replicability may be a result of more stringent filtering of marginally linear barcodes). This problem is expected to be worse on datasets for which exponential growth over the whole length of the experiment cannot be assumed. Therefore, we do not recommend this method when timepoint population sizes are available.

**Score aggregation and data filtering**

We calculate the slope separately for each barcode and timepoint, taking the standard error of the estimate as a starting point for error. Typically, we want to assess scores for protein-level variants, which may be represented by multiple DNA-level variants and/or multiple barcodes. First, we calculate a score for each barcode averaged across replicates. The average score is the arithmetic mean of the slopes, and the error is:

$$\frac{\sqrt{\epsilon_a^2 + \epsilon_b^2 + \ldots \epsilon_n^2}}{n} \quad \{8\}$$

where $\varepsilon_n$ represents the standard error from a single replicate. We then aggregate barcodes that represent the same protein variant. For variants with missense or nonsense mutations, co-occurring synonymous mutations are ignored and all variants that produce the same protein sequence are averaged. Error is propagated as in {8} over n barcodes. We separately calculate the standard error between barcodes that represent the same protein variant. The number of barcodes is also reported for each variant. For synonymous variants, barcodes are aggregated across all variants with synonymous mutations in the same codon and no co-occurring mutations. Variants with synonymous mutations in multiple codons and no missense mutations are discarded. Barcodes are aggregated under the label 'Wild type' if the variant they represent exactly matches the wild type DNA sequence.

We impose data quality filters at four levels. First, barcodes are discarded if they do not meet a minimum threshold for input reads, which we set at 20. We observe that barcodes with fewer than 20 input reads have higher estimated error, on average. Second, if barcodes are significantly non-linear when regressing term {6} over time, we discard it. We set a static threshold at S.E. = 0.2, discarding barcodes above this threshold. Third, we perform a repeated-measures ANOVA for each barcode to detect whether different replicates produce significantly different terms from {6} across timepoints. Barcodes falling below a significance threshold, which we set at p = 0.05, are discarded. Fourth, we discard variants with significant disagreement between synonymous barcodes. If the standard error across all barcodes for a variant exceeds a static threshold, which we set at S.E. = 0.2, we discard the variant.

**Modelling epistasis**

For each double missense variant for which we scored each single missense variant composing it, we estimated the progeny expected for the double missense variant as follows:

$$r_{exp} = \frac{r_1 * r_2}{r_{wt}} \quad \{9\}$$

where $r_{exp}$ is the expected growth rate; $r_1$ and $r_2$ are the measured growth rates of the single missense variants; and $r_{wt}$ is the mean growth rate of variants synonymous to the wild type sequence. We estimated standard error for the expectation as follows:

$$SE_{exp} = \sqrt{SE_1^2 + SE_2^2 + 2SE_{wt}^2} \quad \{10\}$$

We calculated growth rates and error for the double missense variant using the same procedure as for single missense variants. We then compared the empirical growth rate to the expected growth rate for each variant. Variants were significantly epistatic if:

$$t = \frac{r - r_{exp}}{\sqrt{SE^2 + SE_{exp}^2}} > 1.96 \quad \{11\}$$

**Determining promiscuity**

For each single missense variant that we scored on all four receptors, we estimated promiscuity as a rough estimate of the average infectivity over the space of possible receptor, relative to wild type. We calculated an ordinary least squares regression line, y = mx +b, of the variant-specific growth rate on each receptor over the mean growth rate of synonymous variants on each receptor. We calculated the area under the curve for each variant divided by the area under the curve for synonymous variants. For variants with a positive intercept, this was:

$$AUC = \frac{r_{wt}(r_{wt}m + 2b)}{2} \qquad \{12\}$$

While for variants with a negative intercept, this was:

$$AUC = \frac{(r_{wt}m + b)(r_{wt} + b/m)}{2} \qquad \{13\}$$

**Supplementary Figures**

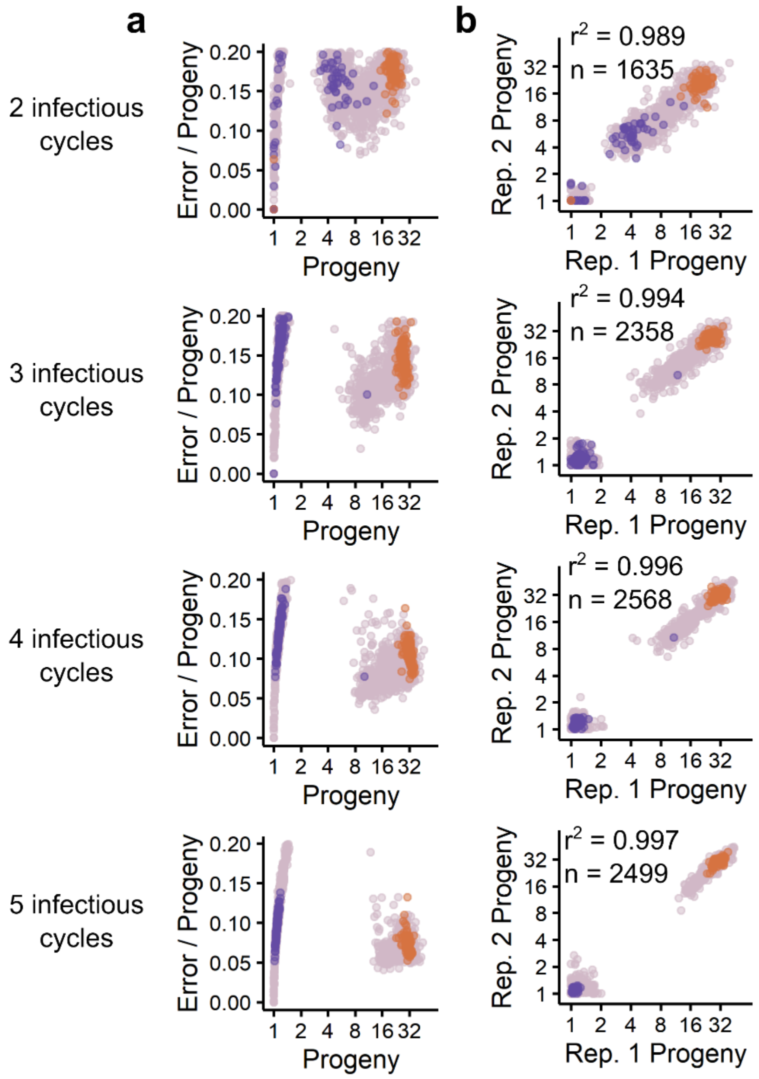

***Figure S1: Replicability of selection for λ infectivity, as assessed by Model-Bounded Scoring.*** *(a) As more infectious cycles are added, the estimated error per variant decreases and the scoring better separates the nonsense and synonymous distributions. However, we start to lose variants that are strongly deleterious but not null-like as they get swept from the population, which is increasing in average fitness over time. We used scores from four infectious cycles in order to balance precision and coverage. Mauve: single missense variants. Orange: synonymous variants. Purple: nonsense variants. (b) Adding infectious cycles improve between-replicate correlation, but only modest improvements are possible.*

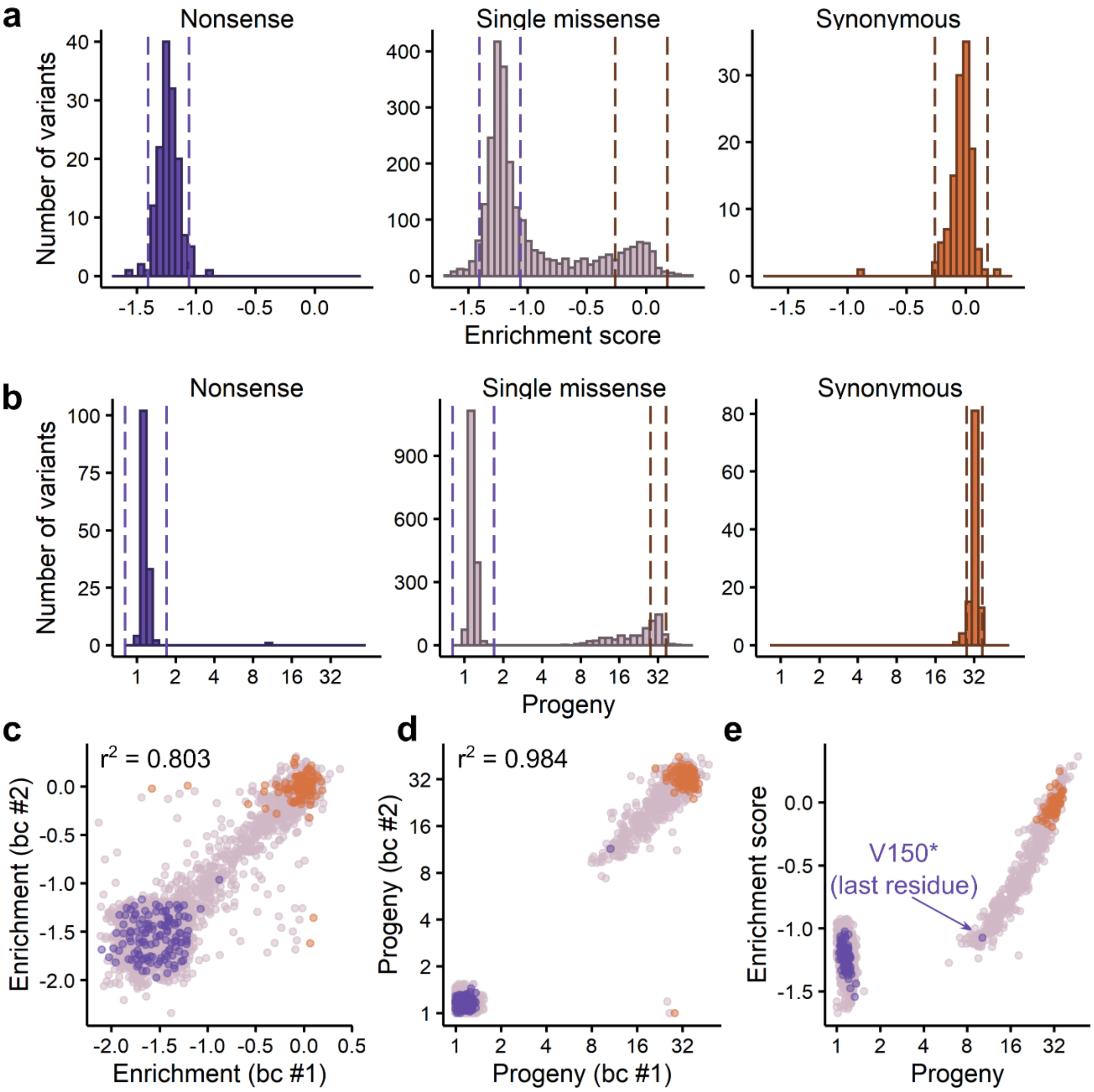


***Figure S2: Comparison of Model-Bounded Scoring with Enrichment-based scoring.*** *(A) The distribution of fitness effects, using enrichment-based scoring. The Enrich2 scoring metric was used, but we separately wrote the code to match filters and score aggregation with our Model-Bounded Scoring approach. (B) The distribution of fitness effects using Model-Bounded Scoring produces a tighter distribution of synonymous variants and better separation of nonsense and synonymous variants than enrichment-based scoring. (C) For our enrichment-based scores, we compare the score for each variant between two randomly selected barcodes. (D) For Model-Bounded Scoring, we compare the progeny for each variant between two randomly selected barcodes. We see generally better replicability*

*than enrichment-based scoring. (E) Directly comparing MBS-derived progeny and enrichment scores shows that MBS is effectively segregating low-fitness variants into those that are rare but measurable and those that are not measured. This segregation likely reflects fitness, because all nonsense variants are put in the null-like category except that at the last position, which might be expected to have non-zero fitness.*

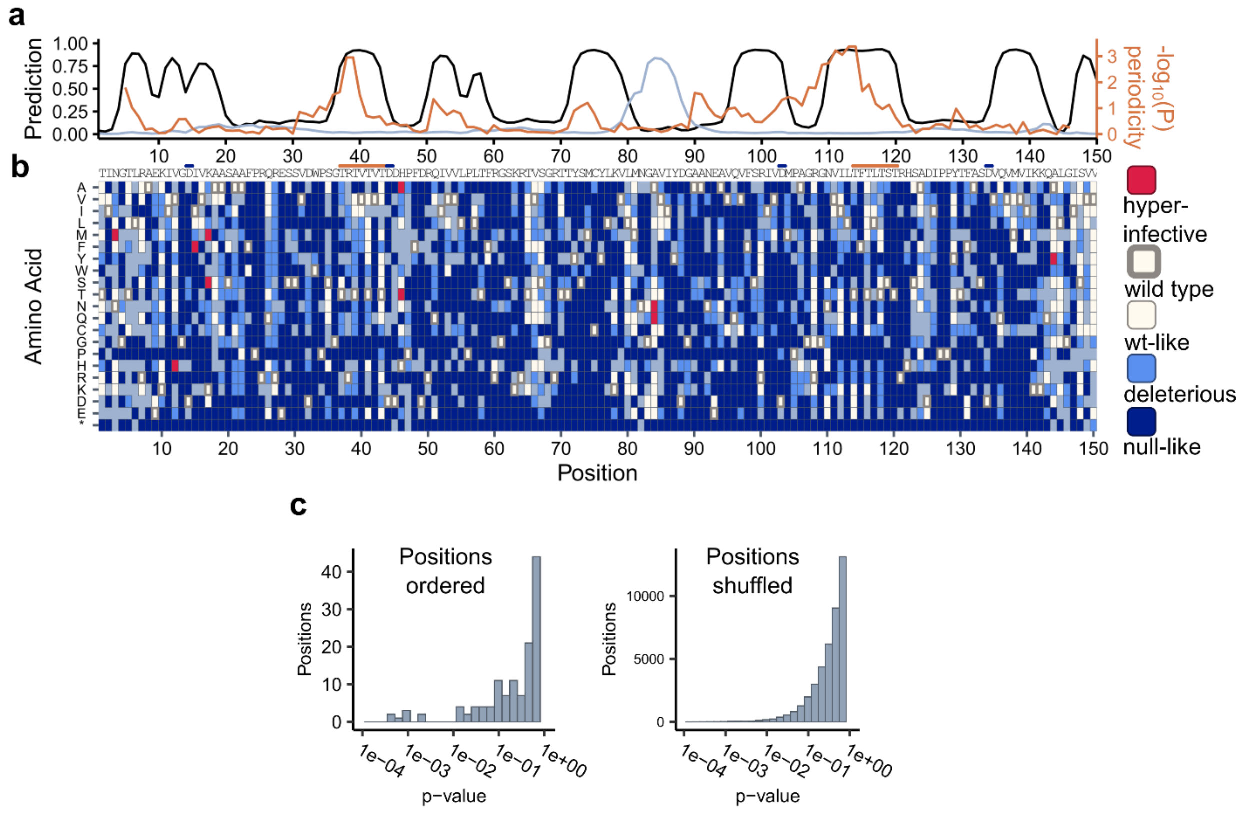


***Figure S3: Patterns of mutational tolerance reflect prevalence of β-sheets. (a)*** *Predictions of β-sheets (black) and α-helices (silver) from Jpred [Drozdetskiy et al. 2015]. Predicted β-sheets strongly overlap with regions of significant periodicity (orange). Periodicity was calculated by running a student's t-test between the average progeny produced my single missense variants in positions in one or the other period (period=2), using a 10-residue sliding window.* ***(b)*** *Heatmap showing the categorical effects of mutations. Glycine and Proline are more disfavored, on average, than other residues.* ***(c)*** *Statistical basis for determining significantly periodic regions. Positions were shuffled repeatedly, and the approach described in (a) was used to detect periodic 10-residue windows. The shuffled positions were very unlikely to yield p-values below $10^{-2}$, whereas several windows below this threshold were present in the properly ordered data.*

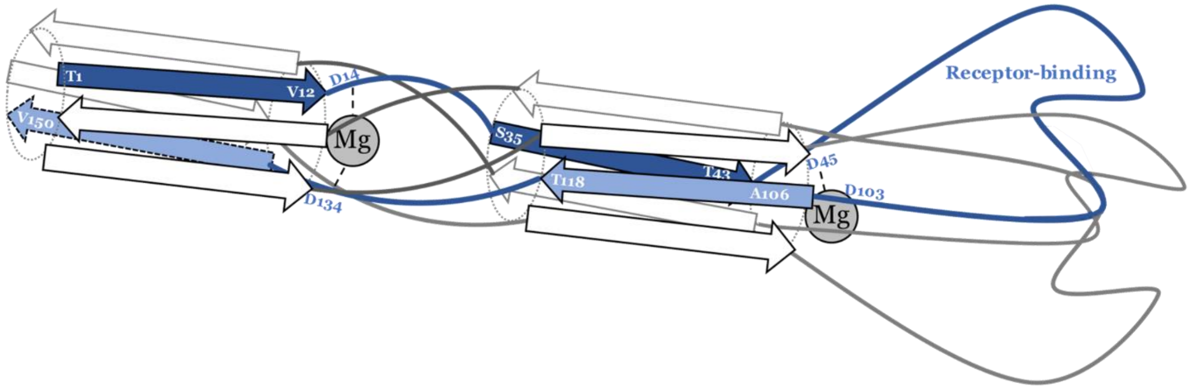


***Figure S4: Coarse structural model for consideration of J structural features.*** *Based on the patterns of mutational tolerance and using the T4 tail fiber structure [Bartual et al. 2010] as a general guide, we propose this coarse model of J's structure. Antiparallel β-sheets form a helical barrel in the 'stalk' of the protein, with junctions between these sheets and less-structured regions defined by Mg-coordinating aspartate residues.*

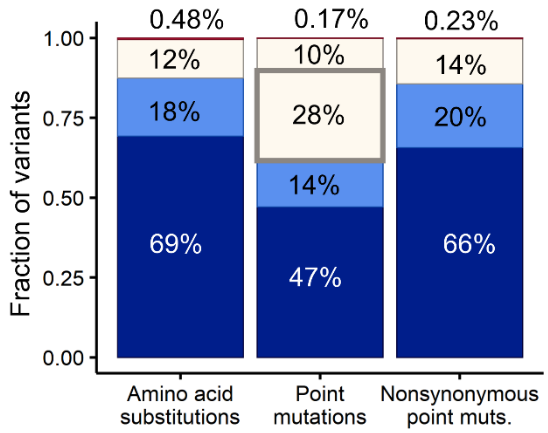


***Figure S5: Relative distributions of categories between all codon substitutions and point mutations only.*** *The fraction of single missense variants that we detected falling into each phenotypic category. Red: hyper-infective variants, white: wt-like variants (thin border) or wild type variants (thick border), light blue: deleterious variants, navy: null-like variants. We calculated fractions for all substitutions, as well as point mutations (including synonymous) and nonsynonymous point mutations. Point mutations include amino acid substitutions possible by point mutations: the actual DNA sequence was not required to have only a single base change.*

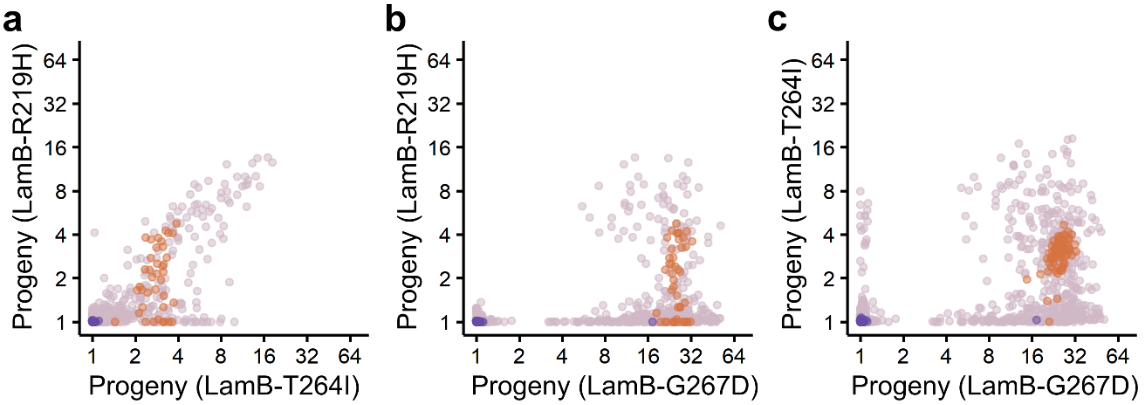


***Figure S7: Comparison of J variant infectivity between multiple novel hosts.*** *For each pairwise comparison of hosts, excluding LamB$^{wt}$, the infectivity of all J variants measured on both hosts is shown.* ***(a)*** *LamB-R219H by LamB-T264I.* ***(b)*** *LamB-R219H by LamB-G267D.* ***(c)*** *LamB-T264I by LamB-G267D. For all hosts, the most infective viral variants are likely to be highly infective on the other host as well.*

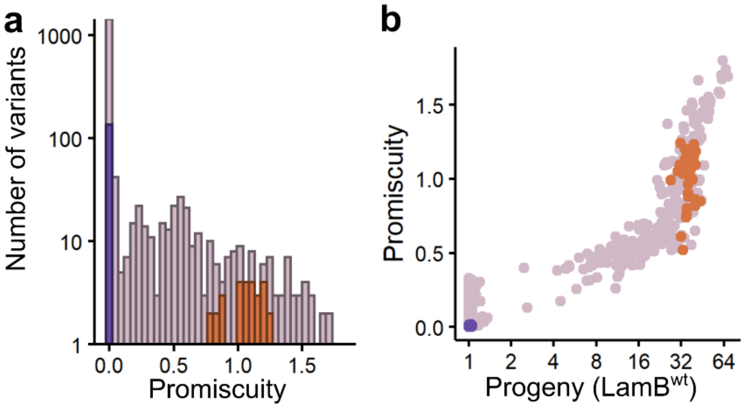


***Figure S8: Distribution of promiscuity****. Promiscuity, as described in Figure 5, is shown for all synonymous (orange), nonsense (purple) and single missense (mauve) variants. (A) The distribution of promiscuity scores. Scores are calculated relative to wild type, such that the mean promiscuity of synonymous variants is 1. (B) Promiscuity is positively, though not linearly, associated with infectivity. The most promiscuous variants are therefore neutral-to-beneficial on a wild type host.*

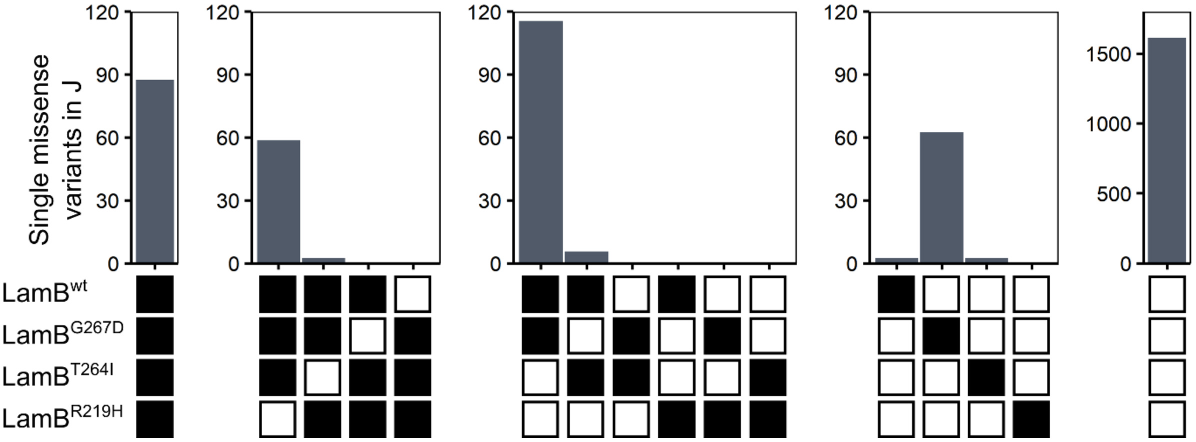


***Figure S9: Subsets of resistance alleles infected by J variants.*** *For each possible combination of hosts (bottom, black: infection, white: no infection), the number of J single missense variants that infect those hosts is shown. The left-most bar of each panel is consistent with a nested model, as it includes infection on only the least resistant alleles. LamB-G267D was more likely to be specifically infected than LamB-wt, perhaps because wild type J has already been selected for specificity to LamB-wt.*

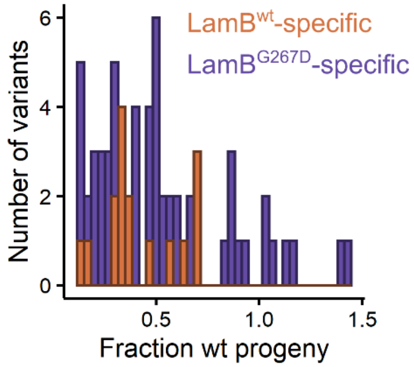


***Figure S10: Distribution of scores for LamB$^{wt}$- and LamB$^{G267D}$-specific variants, on the host they are specific for.*** *For J variants that are specific to LamB$^{G267D}$ or LamB$^{wt}$, the deleteriousness of those variants is shown as their progeny relative to wild type J. LamB$^{wt}$-specific variants are much less likely to be neutral or beneficial. We posit that this bias arises because wild type J has already been selected for specificity to LamB$^{wt}$, removing those variants except when they have a significant cost.*

| | | selection 1 | | | selection 2 | | | | |
|---|---|---|---|---|---|---|---|---|---|
| J variant | *E. coli* allele(s) isolated from | Progeny | Error / estimate | barcodes | Progeny (wt) | Progeny (R219H) | Progeny (T264I) | Progeny (G267D) | Promiscuity |
| S29G | LamB$^{G151D}$ | 1.17 | 0.14 | 6 | not measured | not measured | 1.27 | 1.05 | not measured |
| S30G | LamB$^{-}$ | 1.08 | 0.11 | 20 | 1.00 | 1.00 | 1.00 | 1.02 | 0.00 |
| T58M | LamB$^{S247L}$,LamB$^{G246R}$,LamB$^{-}$ | 1.08 | 0.10 | 11 | 1.02 | 1.03 | 1.00 | 1.00 | 0.01 |
| E93V | LamB$^{G151D}$ | 1.15 | 0.14 | 15 | 1.00 | 1.00 | 1.02 | 1.00 | 0.00 |
| A94S | LamB$^{G151D}$ | 28.73 | 0.10 | 23 | 42.00 | 6.60 | 9.20 | 11.03 | 1.40 |
| V95A | LamB$^{S247L}$ | 1.10 | 0.11 | 29 | 1.01 | 1.00 | 1.00 | 1.01 | 0.00 |
| Q96R | LamB$^{G151D}$ | 1.09 | 0.11 | 58 | 1.01 | 1.00 | 1.00 | 1.01 | 0.00 |
| H122L | LamB$^{-}$ | 1.18 | 0.15 | 12 | 1.06 | 1.00 | 1.01 | 1.00 | 0.01 |
| D125K | LamB$^{-}$ | not measured | not measured | not measured | not measured | not measured | 2.62 | 1.00 | not measured |
| L145P | LamB$^{E148K}$, LamB$^{G245R}$ | 1.10 | 0.12 | 4 | 1.03 | 1.00 | 1.00 | 1.00 | 0.00 |

***Table S1: Previously published host range mutations detected in our assays.*** *For host range mutations reported in Meyer et al. (2012) or Werts et al. (1994), we report the estimated progeny in our selections. Most variants were strongly deleterious in our assays. Most of these host range mutations were identified in the context of other* *mutations, and we did not capture most of the double missense combinations. Therefore, these mutations could be advantageous in the context of positive epistasis in other assays.*
